# Adult Plant Resistance in Maize to Northern Leaf Spot Is a Feature of Partial Loss-of-function Alleles of *Hm1*

**DOI:** 10.1101/244491

**Authors:** Sandeep R. Marla, Kevin Chu, Satya Chintamanani, Dilbag Multani, Antje Klempien, Alyssa DeLeon, Kim Bong-suk, Larry D. Dunkle, Brian P. Dilkes, Gurmukh S. Johal

## Abstract

Adult plant resistance (APR) is an enigmatic phenomenon in which resistance genes are ineffective in protecting seedlings from disease but confer robust resistance at maturity. Maize has multiple cases in which genes confer APR to northern leaf spot, a lethal disease caused by *Cochliobolus carbonum* race 1 (CCR1). The first identified case of APR in maize is encoded by a hypomorphic allele, *Hm1*^*A*^, at the *hm1* locus. In contrast, wild type alleles of *hm1* provide complete protection at all developmental stages and in every part of the maize plant. *Hm1* encodes an NADPH-dependent reductase, which inactivates HC-toxin, a key virulence effector of CCR1. Cloning and characterization of *Hm1*^*A*^ ruled out differential transcription or translation for its APR phenotype and identified an amino acid substitution that reduced HC-toxin reductase (HCTR) activity. The possibility of a causal relationship between the weak nature of *Hm1*^*A*^ and its APR phenotype was confirmed by the generation of two new APR alleles of *Hm1* by mutagenesis. The HCTRs encoded by these new APR alleles had undergone relatively conservative missense changes that partially reduced their enzymatic activity similar to HM1^A^. No difference in accumulation of HCTR was observed between adult and juvenile plants, suggesting that the susceptibility of seedlings derives from a greater need for HCTR activity, not reduced accumulation of the gene product. Conditions and treatments that altered the photosynthetic output of the host had a dramatic effect on resistance imparted by the APR alleles, demonstrating a link between the energetic or metabolic status of the host and disease resistance affected by HC-toxin catabolism by the APR alleles of HCTR.

**AUTHOR SUMMARY:** Adult plant resistance (APR) is a phenomenon in which disease resistance genes are able to confer resistance at the adult stages of the plant but somehow fail to do so at the seedling stages. Despite the widespread occurrence of APR in various plant diseases, the mechanism underlying this trait remains obscure. It is not due to the differential transcription of these genes, and here we show that it is also not due to the differential translation or activity of the APR alleles of the maize *hm1* gene at different stages of development. Using a combination of molecular genetics, biochemistry and physiology, we present multiple lines of evidence that demonstrate that APR is a feature or symptom of weak forms of resistance. While the mature parts of the plant are metabolically robust enough to manifest resistance, seedling tissues are not, leaving them vulnerable to disease. Growth conditions that compromise the photosynthetic output of the plant further deteriorate the ability of the seedlings to protect themselves from pathogens.

**One sentence summary:** Characterization of adult plant resistance in the maize-CCR1 pathosystem reveals a causal link between weak resistance and APR.

## INTRODUCTION

Plant responses to pathogens are dynamic, and they involve a number of inducible mechanisms that are tightly regulated both in space and time (1). They are called into action only at the time and site of infection. The tight regulation of innate immunity is due to disease resistance (DR) genes that plants inherit from their parents and which often segregate with the trait of resistance (1–3). A vast majority of these DR genes function in every part of the plant and at every stage of development. However, many exceptions exist where resistance is manifested in a tissue-or developmental stage-specific manner. In most instances of developmentally regulated resistance, plants are susceptible at the seedling stage but become increasingly resistant toward maturity. The term commonly used to define such developmentally regulated resistance is adult plant resistance (APR), although other terms such as age-associated resistance, ontogenic resistance, mature plant resistance, or flowering-induced resistance have also been used in the literature to describe the same phenomenon (4–8).

Adult plant resistance (APR) often manifests gradually with the advancement of plant age, but a few cases have been reported where the onset is abrupt, happening sharply at a certain stage of development (9–12). An example of the latter kind is the wheat *Lr34* gene-mediated resistance, in which the onset against the leaf rust pathogen, *Puccinia triticina,* is largely confined to the uppermost leaf (flag leaf) (13). In contrast, in the rice-*Xanthomonas oryzae* pv. *oryzae* pathosystem, resistance conferred by the *Xa21* gene is almost negligible during the first three weeks of age but then increases steadily each week, reaching full efficacy at maturity (14,15). Similarly, the *Yr36*-conferred resistance in wheat to *Puccinia striiformis* (16) and the *Hm2*-conferred resistance in maize to *Cochliobolus carbonum* race 1 (CCR1) increase gradually with plant age (12).

In efforts to understand the mechanistic basis of APR, several genes conferring this form of resistance were isolated in different pathosystems. Some of these genes include *Cf-9B* from tomato conferring resistance to leaf mold (17), *Mi-1* from tomato conferring resistance to aphids (18), *Xa21* from rice conferring resistance to leaf blight (14), *Lr67* and *Lr34* from wheat conferring resistance to leaf rust (13,19), *Yr36* from wheat conferring resistance to stripe rust (16), and *Hm2* from maize conferring resistance to leaf blight (12). Two of these genes, *Cf-9* and *Mi-1*, clearly follow the gene-for-gene (GFG) paradigm in conferring resistance, while four others, *Lr67, Lr34, Yr36* and *Hm2,* do not, suggesting that any disease resistance gene has the potential to confer an APR phenotype.

What makes a gene behave in an APR manner? This question still eludes us, even though a number of APR genes, including those described in the preceding paragraph, have been cloned and characterized. One logical expectation was that the phenotype of APR genes may derive from their differential expression at different stages of plant development and that the level of gene expression would match their phenotypic efficacy closely. However, this has been ruled out with the majority of the APR genes, as their transcript levels do not reflect changes in their resistance phenotype (12,13,15,17,20). Other possibilities that may affect the APR behavior of these genes are differential translation, differential post-translational modifications, and developmental changes in plant physiology and metabolism.

To gain insight into the mechanistic basis of APR in maize, we have been studying the northern leaf spot (NLS) disease of maize (*Zea mays*) caused by *C. carbonum* race 1. A classic APR syndrome is described in this pathosystem where alleles at two homeologous loci can confer resistance in a developmentally programmed fashion (9). These duplicate genes, *Hm1* and *Hm2*, encode NADPH-dependent HC-toxin reductases (HCTR), which utilize NADPH as a cofactor to reduce an essential ketone function in HC-toxin (HCT), the key disease causing effector of CCR1, and abolish its activity (21–23). There is one prominent difference between the HCTRs encoded by *hm1* and *hm2*: whereas the HCTR encoded by wild type (WT) *Hm1* contains 356 amino acids, the HCTR encoded by the functional *Hm2* allele is truncated and lacks the last 52 amino acids compared to HM1 (12). This truncated allele is the only functional allele that has been identified at *hm2*, and it confers APR against CCR1 when *hm1* is null. *Hm2* is expressed throughout the age of the plant (12), ruling out developmentally regulated transcript accumulation as the mechanism of APR. Like *Hm2*, an allele of *hm1* conferring APR has also been described (9). Designated *Hm1*^*A*^, this APR allele is recessive to the WT *Hm1* allele and dominant to the *hm1* null allele (9).

To explore why and how the *Hm1*^*A*^ allele leads to an APR phenotype, we have cloned and characterized it in detail. Comparison of the sequence of *Hm1*^*A*^ with those of the WT haplotypes from a number of resistant inbreds and accessions revealed a single amino acid substitution in the HM1^A^ peptide that is unique to its APR behavior. HM1^A^ transcripts accumulated to similar levels throughout plant growth and development, as did the translational product of the gene. However, the HCTR activity in *Hm1*^*A*^ plants was intermediate between WT (*Hm1Hm*1) and null mutant (*hm1hm1hm2hm2*) plants. This, along with the truncated nature of the APR allele at *hm2*, prompted us to consider if the hypomorphic *Hm1* allele in *Hm1*^*A*^ was the reason for its APR phenotype. This hypothesis was addressed by mutagenesis, generating two new APR mutants of the B73 maize inbred, which is homozygous for the WT allele at *hm1* and the null allele at *hm2* (*Hm1Hm1hm2hm2*). Both new APR alleles were found to contain single amino acid substitutions in HM1-B73 and reduced HCTR activity. Thus, APR is a symptom of partial loss-of-function mutations in *Hm1* that result in seedling susceptibility.

## RESULTS

### Detailed genetics of APR-conferring *Hm1*^*A*^ as an allele of *hm1*

The APR trait attributed to *Hm1*^*A*^ was first noticed in the inbred P8, developed at Purdue University in the early 1960s (9). The genetic evidence linking the APR of P8 with an allele of *hm1* (*Hm1*^*A*^) made use of two segregating populations, a testcross and an F_2_ population, generated by crossing P8 (*Hm1*^*A*^*Hm1*^*A*^*hm2hm2*) with the resistant inbred WF9 (*Hm1Hm1Hm2Hm2*). The susceptible inbred for the testcross was Pr, which is homozygous for null mutations at both the *hm1* and *hm2* loci. There were at least two concerns with this study. First, it used a relatively small number of progenies, comprising about 90 plants each for both the F_2_ and testcross populations. Second, the resistant inbred WF9 also contained an APR allele at the *hm2* locus, leaving room for error in extrapolation from these data.

These concerns necessitated that we revisit these findings, to clone and characterize *Hm1*^*A*^. We acquired P8 from the Germplasm Resources Information Network (GRIN). To confirm that this source of P8 harbored the *Hm1*^*A*^ allele reported by Nelson and Ullstrup (1964), we conducted a thorough analysis of the genetics of P8 resistance to CCR1. We first crossed P8 twice with Pr (*hm1hm1hm2hm2*) to produce a BC_1_F_1_ testcross population. Of 384 BC_1_F_1_ plants inoculated with CCR1, 186 plants were susceptible at both the seedling and adult stage while 198 plants were susceptible as seedlings, but later emerging leaves were fully resistant, consistent with the APR phenotype of P8. The recessive null *hm1* allele of Pr (designated as *hm1*^*Pr*^) contains a 256-bp *Drone* transposon insertion in exon 4 (24). All 186 plants susceptible at maturity were homozygous for *hm1*^*Pr*^, whereas all 198 plants that were initially susceptible and then displayed APR were heterozygous for *hm1*^*Pr*^. This 1:1 ratio of susceptible *vs.* APR plants (X^2^ −0.375, *P* > 0.05, 1 d.f.) indicated that a single gene at or near the *hm1* locus controlled the APR behavior of P8.

Next we crossed P8 to Pr1, a near isogenic line (NIL) of Pr in which the mutant *hm1* allele was replaced by a WT *Hm1* (25). The resulting *Hm1*^*A*^*Hm1* F_1_ hybrid was testcrossed to Pr, the *hm1hm2* null stock. The inheritance of *Hm1*^*Pr*1^ vs. *Hm1*^*A*^ in this population was tracked with a PCR-based marker that differentiated between those two alleles. Of the 540 F_1_ test cross progeny, 276 were susceptible as seedlings and later exhibited APR, while the remaining 264 were completely resistant to CCR1 regardless of age. All 264 completely resistant plants had inherited the WT *Hm1* allele from Pr1, while the 274 plants that exhibited APR had inherited the *Hm1*^*A*^ allele from P8. Chi-squared tests supported the 1:1 expected inheritance of monogenic inheritance (X^2^ −0.266667, *P* > 0.05, 1 d.f.). No recombinants between the genotypes at the *hm1* locus and the expression of CCR1 susceptibility were found in either population (924 opportunities for crossover). This confirmed that the source of P8 we obtained recapitulated the phenomenon described in 1964 (9) and that the APR of P8 is likely conferred by the *Hm1*^*A*^ allele.

To incorporate *Hm1*^*A*^ into a uniform background for detailed phenotypic comparisons, we introgressed this APR allele into the B73 inbred by crossing P8 (*Hm1*^*A*^*Hm1*^*A*^ *hm2hm2*) to B73 (*Hm1Hm1hm2hm2*). As *Hm1*^*A*^ is recessive to WT *Hm1*, we utilized sequence polymorphism between *Hm1*^*A*^ and *Hm1*^*B73*^ to construct a PCR-based marker. After seven crosses to B73 with selection for the *Hm1*^*A*^ genotype, BC_7_F_2_ progeny were generated by self-pollinating a heterozygous plant. This BC_7_F_2_ population segregated in a 3:1 ratio for complete resistance and APR, again consistent with *Hm1*^*A*^ being responsible for APR of P8. Homozygous *Hm1*^*A*^ plants from this population were selected and maintained as an *Hm1*^*A*^ near-isogenic line in B73.

### Phenotypic manifestation of adult plant resistance in maize to CCR1

To develop a comprehensive account of the onset of APR by *Hm1*^*A*^, we also introgressed the null *hm1*^*Pr*^ allele into the B73 background over seven generations, and crossed with *Hm1*^*A*^ B73 NIL to generate plants heterozygous for *Hm1*^*A*^. Both homozygous (*Hm1*^*A*^*Hm1*^*A*^) and heterozygous (*Hm1*^*A*^*hm1*^*Pr*^) *Hm1*^*A*^ plants were inoculated with CCR1 at weekly intervals, starting at 1 week-after-planting (wap) and culminating at 10 wap. Their infection phenotypes were measured using a 1-10 disease rating scale (12) and compared with those of B73 and a B73 NIL containing the null *hm1* allele (*hm1*^*Pr*^ B73 NIL). A rating of 10 on this scale indicated highly susceptible plants, while a rating of 1 indicated complete resistance.

The susceptible *hm1*^*Pr*^ B73 NILs scored 10 on the disease rating scale regardless of age, and the resistant controls (B73 inbred), which produced small chlorotic flecks in response to CCR1 infection, scored 1 throughout development. Plants containing *Hm1*^*A*^ exhibited very little resistance at the seedling stage, but severity scores decreased with age (Fig 1A and 1C). At the age of week-1, *Hm1*^*A*^ seedlings were consistently rated 8 or higher. This disease rating dropped to 5 or less by week-5. At week-10, *Hm1*^*A*^ plants resembled the resistant controls, receiving a rating of 1 (Fig 1B and 1C). The level of resistance conferred by *Hm1*^*A*^ correlated with the age of the whole plant at the time of inoculation and not the age of the inoculated leaf. Inoculating each leaf of *Hm1*^*A*^*Hm1*^*A*^ and *hm1hm1* plants at week-5 of plant growth confirmed this observation. All the leaves of *Hm1*^*A*^ plants were equally resistant regardless of their age, and all the leaves of *hm1hm1* plants were equally susceptible (data not shown).

**Figure 1.**
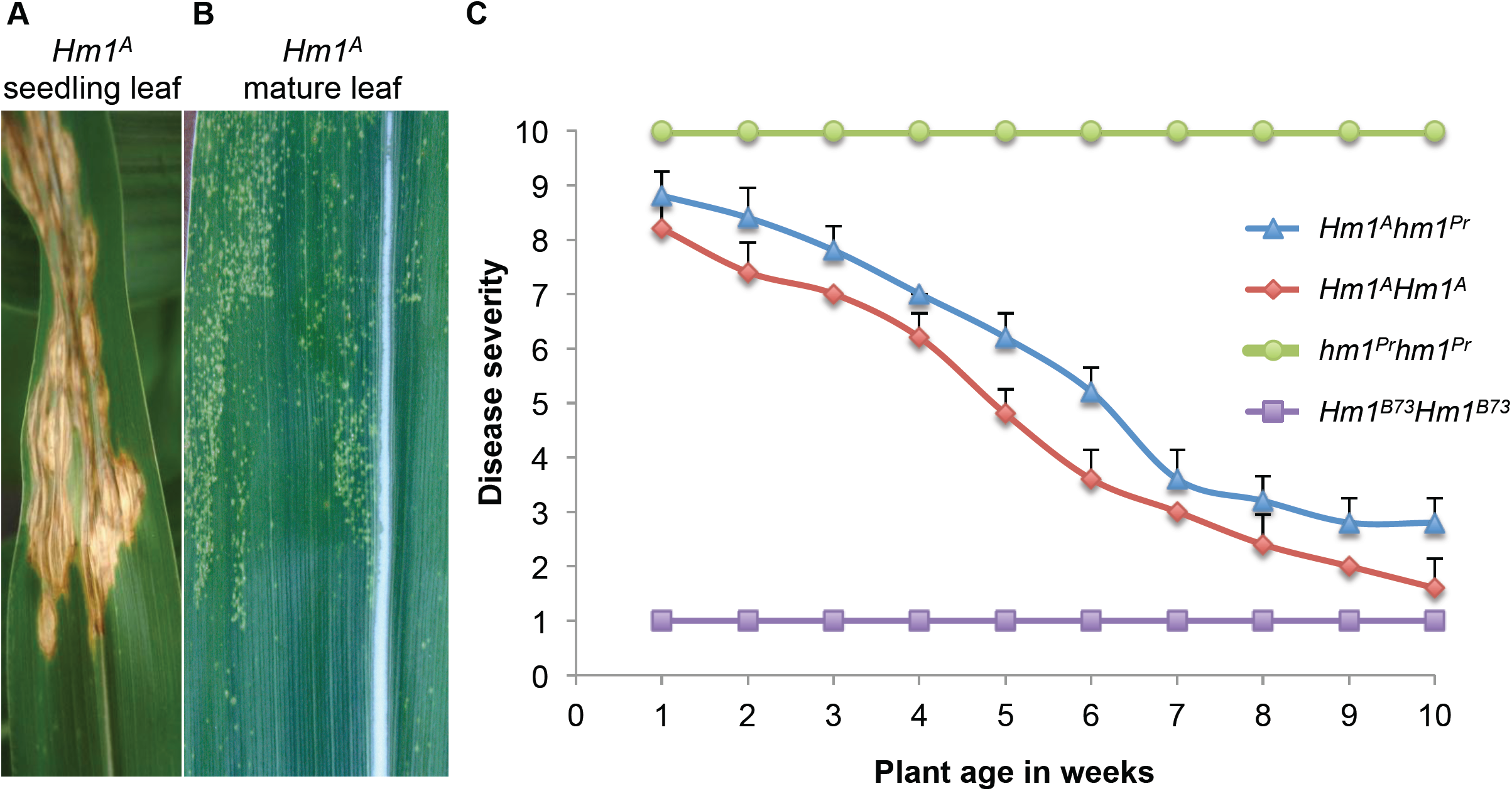
Developmental onset of the adult plant resistance phenotype of *Hm1*^*A*^. **(A)** A seedling *Hm1*^*A*^ leaf exhibiting susceptibility to *Cochliobolus carbonum* race 1 (CCR1) at the 2-week age. **(B)** A 9-week old *Hm1*^*A*^ leaf completely resistant to CCR1. **(C)** The disease/resistance phenotype of *Hm1*^*A*^ plants homozygous and heterozygous (*Hm1*^*A*^*hm1*^*Pr*^) for the APR allele to CCR1 at weekly intervals from week-1 through week-10. Ratings were established by controls *Hm1*^*B73*^*Hm1*^*B73*^ (rated 1 and resistant throughout) and *hm1*^*Pr*^*hm1*^*Pr*^ (rated 10 and susceptible throughout). All *hm1* alleles were in the B73 genetic background. Error bars represent standard error calculated using R statistical package.

Similar to the APR conferred by the *Hm2* gene (12), the resistance conferred by *Hm1*^*A*^ was dosage dependent. Plants homozygous for *Hm1*^*A*^ were slightly more resistant to CCR1 at almost all stages of development compared to plants heterozygous for *Hm1*^*A*^ and the null allele (*Hm1*^*A*^*hm1*^*Pr*^) indicating that *Hm1*^*A*^ is haploinsufficient (Fig 1C). The dosage effect was more pronounced at week-5 and declined after week-7 as the plants matured and became completely resistant.

### Molecular characterization of the *Hm1*^*A*^ allele

Atypical behavior of a disease resistance gene can sometimes result from complex structural changes at the locus, such as an increase in the copy number of the gene or a part of the gene (26,27). To address if such a genetic mechanism also led to the *Hm1*^*A*^ APR, we conducted a Southern blot analysis with P8 DNA digested with a variety of restriction enzymes. Consistent with the genetic data, a single *BamHI* restriction fragment hybridized to *Hm1*-specific probes on these blots, indicating that *Hm1*^*A*^ was a single copy gene in the P8 inbred and that the entire gene was present on a 13 kb restriction fragment (Fig 2A). To clone the *Hm1*^*A*^ gene, a lambda library was constructed from the *Bam*H1-digested P8 DNA restriction fragments migrating on a gel as 12 to 15 kb fragments. We identified and sequenced a clone containing the 13 kb *hm1*-encoding fragment. Sequence analysis indicated that our clone contained the entire coding region of the *Hm1* gene, as well as 3.8 kb of the promoter region.

**Figure 2.**
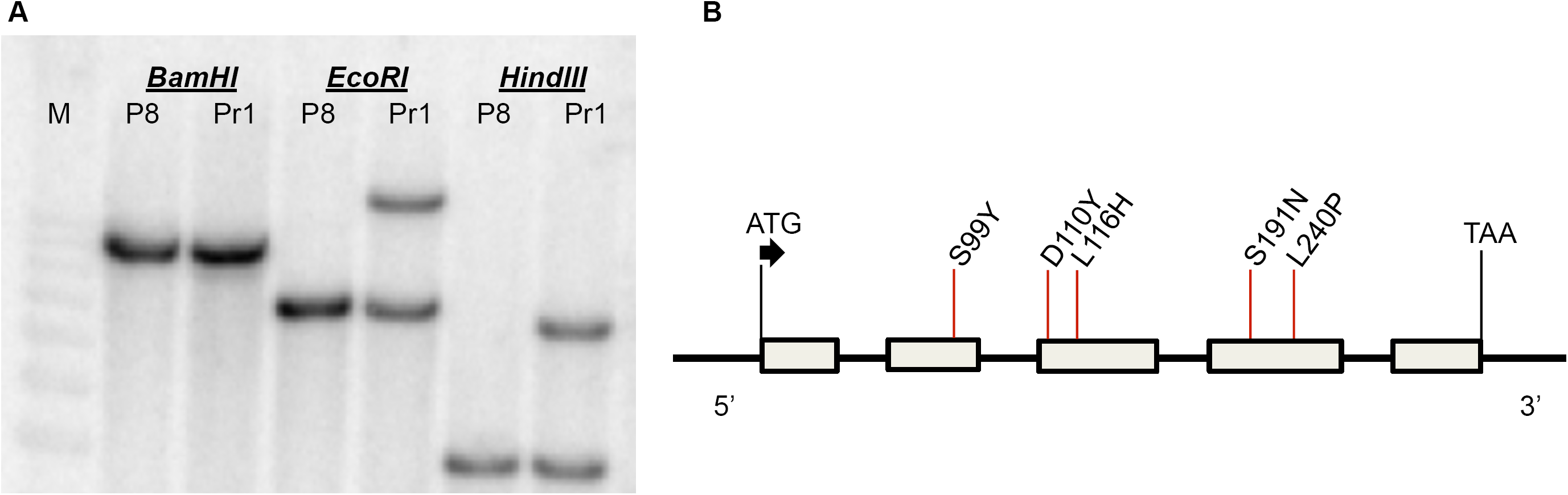
Molecular characteristics of *Hm1*^*A*^. **(A)** Southern blot analysis of DNA of inbreds P8 (*Hm1*^*A*^*Hm1*^*A*^) and Pr1 (*Hm1*^*Pr*1^*Hm1*^*Pr1*^) demonstrating that *Hm1*^*A*^ is a single copy gene. Sample genotypes (inbreds P8 or Pr1) are indicated below the restriction endonuclease used for DNA digestions (BamHI, EcoRI, or HindIII) and M corresponds to the the DNA marker lane. **(B)** Schematic representation of the gene structure of *Hm1*^*A*^ comprised by five exons (grey boxes) and four introns, identical to *Hm1*^*B73*^. The locations and the nature of five amino acids that differ between HM1^A^ and HM1^B73^ are indicated by red lines. The locations of the start and termination codons are also indicated.

To determine the structural changes in *Hm1*^*A*^, its sequence was compared with that of the B73 reference sequence. Significant changes were encountered in the promoter regions of *Hm1*^*A*^ and *Hm1*^*B73*^. Except for a few indels and SNPs, the first −200 bp from the translation start site of the promoter region are similar in *Hm1*^*A*^ and B73 (Fig S1). The next −1.5 kb region upstream, however, is completely different between the two alleles, though this does not seem to be due to the insertion of a transposable element. Interestingly, the promoter region of *Hm1*^*A*^ is identical to that of *hm1*^*Pr*^, the null *hm1* allele from the susceptible inbred Pr. To examine if any other resistant lines containing a wild type *Hm1* allele also had a promoter region identical to that of *Hm1*^*A*^, we used a primer pair designed from the *Hm1*^*A*^ promoter region to PCR amplify DNA from a number of resistant inbreds. Two inbreds, Pr1 and Va35, were found whose *Hm1* WT alleles have the promoter regions identical to that of *Hm1*^*A*^ (Fig S1). Taken together, these results indicate that the promoter polymorphism between *Hm1*^*A*^ and *Hm1*^*B73*^ predicted neither resistance nor susceptibility and thus may be inconsequential to the APR phenotype of *Hm1*^*A*^.

The coding region of *Hm1*^*A*^ also differed from that of *Hm1*^*B73*^, containing nine SNPs. Although four of these SNPs were silent or synonymous, five led to amino acid substitutions in the predicted HM1^A^ peptide (Fig 2B). Relative to the B73 HM1 reference, these substitutions were: a Serine to Tyrosine change at residue 99 (S99Y), an Aspartic acid to Tyrosine change at residue 110 (D110Y), a Leucine to Histidine change at residue 116 (L116H), a Serine to Asparagine change at residue 191 (S191N), and a Leucine to Proline change at residue 240 (L240P) (Fig 2B).

### The L116H substitution is the likely causative polymorphism in the *Hm1*^*A*^ allele

As *Hm1* is one of the most polymorphic genes in maize (28), we decided to examine the peptide sequence of various resistance alleles to potentially pinpoint the amino acid change(s) responsible for the APR behavior of *Hm1*^*A*^. We first amplified and evaluated the HM1 sequences of Pr1 and Va35, the two resistant inbreds that share their promoters with *Hm1*^*A*^, and compared them with the sequences of both HM1^A^ and HM1^B73^. HM1^Pr1^ was found to differ by five amino acids from HM1^B73^, with two of these polymorphisms, S99Y and L240P, also being present in HM1^A^ (Fig S2). These same two changes were also found in HM1^Va35^, which differed from HM1^B73^ by six amino acids. Another resistant *Hm1* allele that differed from B73 by six amino acids was in the inbred W22, but none of those changes matched those of HM1^A^. However, the predicted HM1 of the landrace Enano from Bolivia (28) shared with HM1^A^ the two polymorphisms D110Y and S191N. And most importantly, the HM1 of the landrace Pira from Colombia (28) shared four of the five amino acid changes between HM1^A^ and HM1^B73^. These are S99Y, D110Y, S191N, and L240P, thereby leaving only the L116H polymorphism unique to HM1^A^.

To examine the functional status of the *Hm1* allele of Pira, we acquired this landrace from GRIN and inoculated it with CCR1. It was found to be completely resistant to CCR1, even at the seedling stage. This demonstrated that despite having four of the five amino acid changes of HM1^A^, the *Hm1*^*Pira*^ allele is fully functional and not APR. These results highlight the importance of the L116H substitution in defining the phenotype of *Hm1*^*A*^. Consistent with this hypothesis, the Leucine at 116 is highly conserved not only in all the homoeologs and orthologs of the *Hm1* gene across the grass lineage, but also in the maize dihydroflavonol 4-reductase (DFR), an NADPH-dependent enzyme of the anthocyanin pathway predicted to be a progenitor of HM1 (Fig S2). All these findings suggest that the HM1^A^ L116H substitution is unique to *Hm1*^*A*^ and may underlie its APR behavior to CCR1 in maize by somehow negatively impacting HCTR activity.

### HM1 transcript accumulation is not developmentally regulated in *Hm1*^*A*^

To examine if the transcriptional activity of *Hm1*^*A*^ undergoes any change during plant development, reverse transcription (RT)-PCR was conducted on RNA extracted from CCR1-inoculated *Hm1*^*A*^ plants of diverse ages. Using a semi-quantitative form of this assay, no dramatic changes could be observed in the level of the *Hm1*^*A*^ transcript between the seedling and mature-plant stages (Fig 3A). Likewise, quantitative real time PCR (qRT-PCR) measurements of transcript abundance of *Hm1*^*A*^ plants inoculated with CCR1 at different ages did not detect any rise in HM1 expression as the susceptible plants became resistant over time (Fig 3B). These results ruled out the differential transcription of the *Hm1*^*A*^ allele as the basis for its APR phenotype.

**Figure 3.**
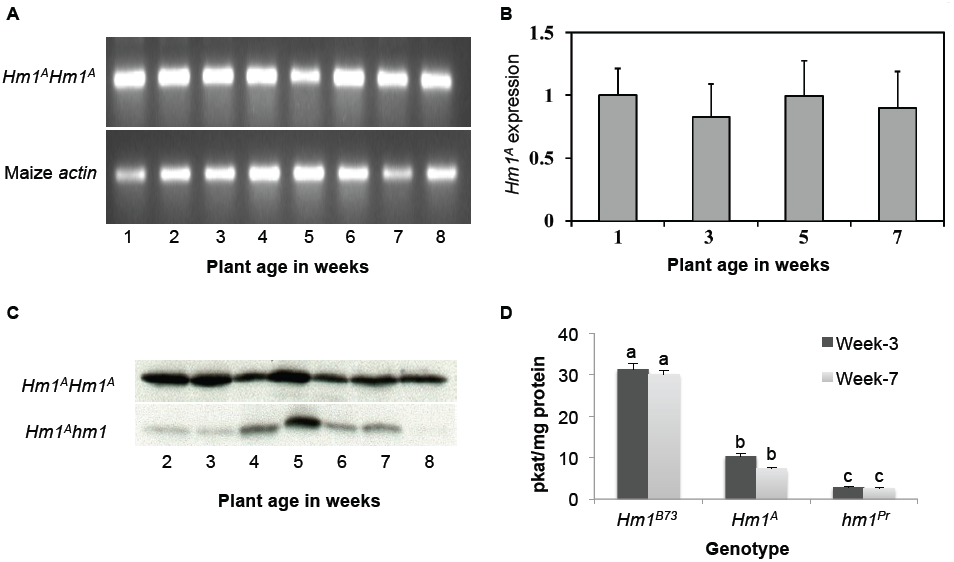
Transcriptional and translational activities of *Hm1*^*A*^ during the seedling and mature stages. **(A)** Reverse transcription (RT)-PCR assay showing no change in *Hm1*^*A*^ accumulation in leaves from week-1 through week-8 after planting. The *actin* gene was used as a control. **(B)** Quantitative real time PCR (qRT-PCR) measurements of the expression of *Hm1*^*A*^ also demonstrates no change in *Hm1*^*A*^ accumulation across the time period when APR is established. **(C)** Western blots showing that the level or stability of the HM1^A^ protein do not change over time during plant development. Equal amounts of protein were loaded following quantification with the Bradford method. **(D)** *In vitro* HC-toxin reductase (HCTR) assays showing that the relative enzymatic activity encoded by *Hm1*^*A*^ is less than *Hm1*^*B73*^ but higher than *hm1*^*Pr*^, the null allele. The specific activity of HCTR varies between alleles but not over time between weeks 3 and 7 in any genotype. The HCTR assay was based on the determination via LC-MS/MS of the amount of HC-toxin reduced by leaf protein extracts from the leaves of all genotypes. Different letters indicate significant differences between genotypes (p_adj_ < 0.05).

### The level and activity of *Hm1*^*A*^-encoded HCTR stays the same during plant development

To address if the differential translational activity of *Hm1*^*A*^ had any impact on its APR behavior, we first conducted western analysis to examine the level and stability of the HM1^A^protein. While an antibody raised against the entire HM1^A^ peptide lacked specificity, a multiple antigenic peptide (MAP) (29) antibody generated against a 13 aa peptide corresponding to residues 312 to 324 of the HM1 peptide worked well and reacted to a single product on Western blots generated from *Hm1*^*A*^ homozygous or heterozygous plants (Fig 3C). No change in the level of the HM1^A^ protein could be detected over time, indicating that the APR phenotype of *Hm1*^*A*^ is not due to differential translation either.

We next addressed if the activity of HCTR encoded by *Hm1*^*A*^ had any role in its APR behavior. To do this, an LC-MS/MS-based *in vitro* HCTR activity assay that quantified the reduction of HC-toxin by crude protein extracts was developed. The *in vitro* measurements were normalized to total protein content, allowing us to estimate the level of the functional HCTR in plant tissues. To examine the level of HCTR over time, proteins were extracted from CCR1-inoculated leaves of 3-and 7-week-old plants of *Hm1*^*A*^ and control stocks, and their HCTR activity was measured in replicated samples. Two trends were noted as shown in Fig 3D. First, the HCTR activity encoded by HM1^A^ was lower than by the WT allele but not null as that of *hm1*^*Pr*^. At both stages of development, the HCTR activity of HM1^A^ was about 3-fold lower than that of HM1. Second, the level of active HCTR differed little if any in 3-or 7-week *Hm1*^*A*^ plants (Fig 3D). Likewise, the HCTR activity of the WT allele also did not differ between week-3 and week-7-old plants (Fig 3D). Two conclusions can be drawn from these results. First, *Hm1*^*A*^ encodes an HCTR that is relatively weaker than the enzyme encoded by the WT allele. Second, the level of the active HCTR stays constant over development and does not account for the APR phenotype of *Hm1*^*A*^.

### Partial loss-of-function mutations confer adult plant resistance in the maize-CCR1 pathosystem

What aspect of the *Hm1*^*A*^ gene structure or function restricts it to be an APR gene, i.e., conferring resistance only at the mature-plant stage but not the seedling stage? Having ruled out differential transcription or translation as possible mechanisms, we paid attention to an attribute of *Hm1*^*A*^ that differentiates it from both the WT and null mutant alleles of *hm1* -the relatively weak nature of the HCTR activity encoded by *Hm1*^*A*^. This partial enzymatic activity of HM1^A^ mirrored exactly the phenotypic strength of resistance conferred by this APR allele, which is recessive to that of WT *Hm1* but dominant to that of null *hm1*. Given that the APR allele at the *hm2* locus also confers partial resistance to CCR1 (12), we pondered if this could be a requirement for a resistance gene to have an APR phenotype.

If this hypothesis that a *Hm1* APR allele owes its phenotype to being a weak or partial loss-of-function allele is correct, we should be able to confirm it by generating new APR alleles from the WT *Hm1* allele by mutagenesis. To address this possibility, we first tried a random mutagenesis screen to generate new alleles of *Hm1*, in large part because of the lethal nature of CCR1 infection on field-grown plants lacking functional *Hm1*. About 1,000 M_2_ families of B73 were generated by treating pollen with the mutagen ethyl methanesulfonate (EMS). Twenty-four plants per M_2_ family were planted in a field and inoculated with CCR1 at the seedling stage. One M_2_ family was identified in which CCR1-susceptible plants segregated in a recessive fashion. These plants remained susceptible throughout their growth, suggesting they were the result of a null mutation. Sequence analysis of the *hm1* allele from this mutant (named *hm1-2*) confirmed its null status and revealed a single G to A transition at the junction of exon3/intron3 as the cause of mutation (Fig S3). Since this change is expected to abolish the splicing of intron 3, it would result in a truncated protein lacking all the amino acids encoded by exons 4 and 5 (Fig S3). It is unlikely that such a grossly truncated protein would have any HCTR activity, and as shown in Fig 6, *hm1-2* exhibited very little enzymatic activity.

We next conducted a targeted mutagenesis screen to generate a series of mutant alleles of *Hm1*. To accomplish this, EMS-mutagenized *Hm1*^*B73*^ pollen was applied to ears of completely susceptible Pr plants in a greenhouse (Fig 4). Approximately 4,500 M_1_ seeds obtained from this cross were planted in the field and inoculated with CCR1 at week-2. Seven plants were identified as CCR1 susceptible at this seedling stage. When inoculated again at week-5, five of them were still fully susceptible, suggesting they were null mutants. The other two plants however exhibited APR as they developed different levels of resistance (Fig S4). Sequencing the *Hm1* gene (Fig S5) from all seven mutants revealed that they all carried GC to AT transitions in the coding region of *Hm1*. The two APR-exhibiting alleles (designated *Hm1-3* and *Hm1-4*) had missense mutations resulting in single amino acid substitutions, T90M in *Hm1-3* and V210M in *Hm1-4*, in the HM1 peptide (Table 1). Of the five null mutants, three (named *hm1-6* to *hm1-8*) had nonsense mutations, one a C82Y substitution (*hm1-5*), and one a splice-site mutation (*hm1-9*) at the junction of intron 4/exon 5 that also produced a pre-mature stop codon (Table 1). The new APR alleles were introgressed back into B73 for seven generations using CAPs markers. Comparison of the resistance phenotype of the two new APR alleles with *Hm1*^*A*^ revealed that all three APR alleles differ markedly from each other in this trait. *Hm1*-3 confers the highest level of resistance at all stages of development, followed by *Hm1*^*A*^ and *Hm1-4* (Fig 5). This screen thus provided us with a series of APR alleles at the *hm1* locus.

**Figure 4.**
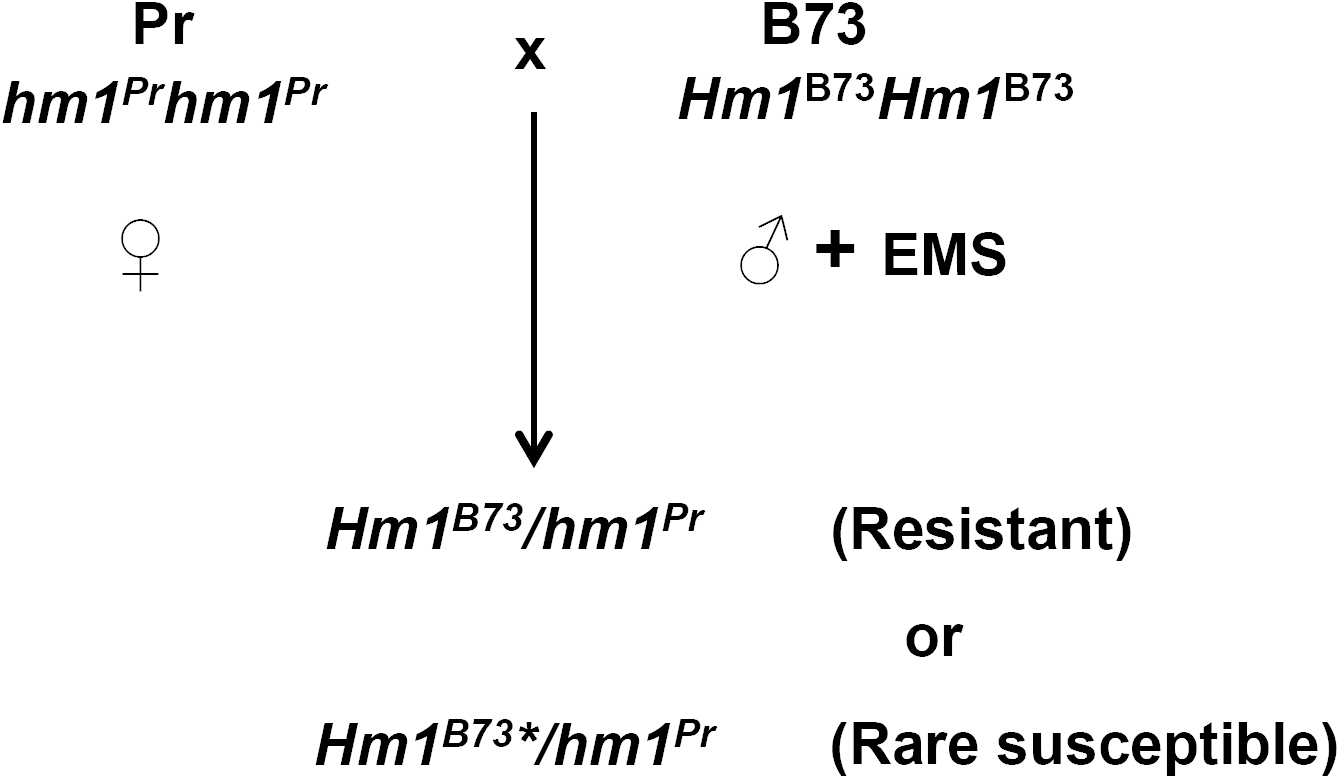
Design of the targeted EMS mutagenesis screen to generate new mutant alleles of *Hm1*. Pollen collected from the fully resistant inbred B73 (*Hm1*^*B*73^*Hm1*^*B73*^) was treated with ethyl methanesulfonate (EMS) and used to pollinate ears of the fully susceptible inbred Pr (*hm1*^*Pr*^*hm1*^*Pr*^) in a greenhouse. The resultant M1 seeds (*Hm1*^*B73*^/*hm1*^*Pr*^) were planted in the field, inoculated with CCR1, and screened for disease resistance at both the seedling stage and at maturity to identify rare susceptible mutants, designated as *Hm1*^*B73*^*\**/*hm1*^*Pr*^. M_1_ mutants that were susceptible at the seedling stage that became resistant with the progression of age were considered APR. Out of about 4,500 M_1_ plants screened, 7 susceptible mutants were found and two became resistant at maturity.

**Figure 5.**
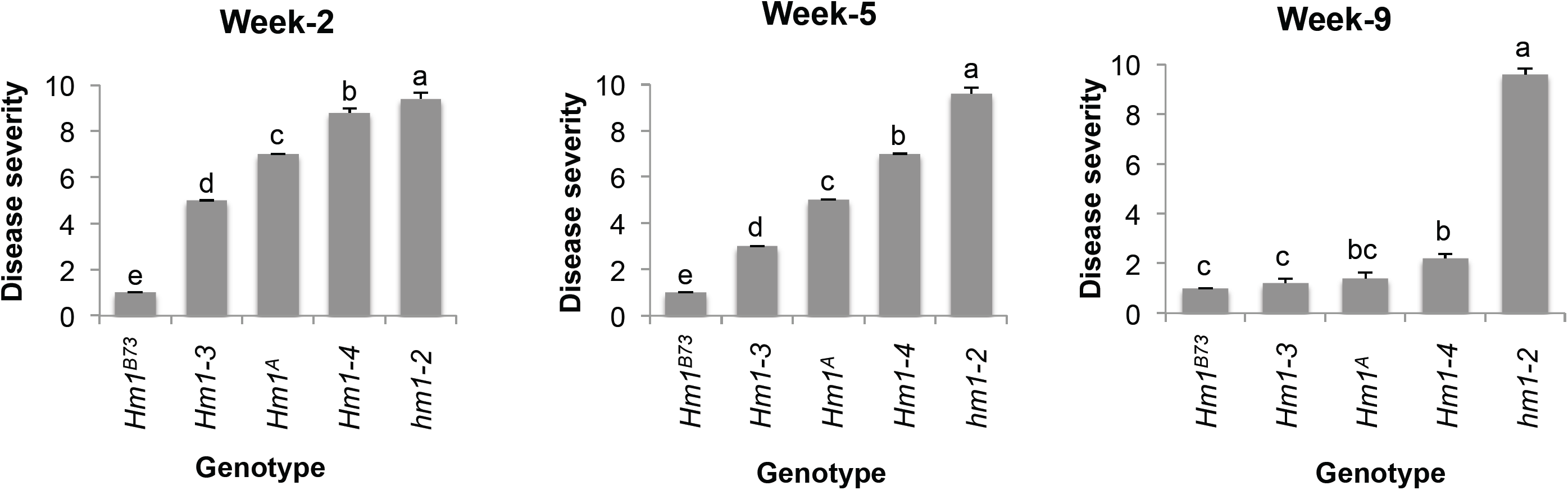
Relative strength of the three APR alleles of *hm1* in conferring protection against CCR1. Like *Hm1*^*A*^, both new APR alleles (*Hm1*-*3 and Hm1*-*4*) were introgressed into B73 for six generations for comparison of their resistance phenotypes. Plants homozygous for the *Hm1*^B73^ and *hm1-2* alleles were fully resistant and susceptible, respectively. Disease resistance evaluations were done three times, at week-2, week-5 and week-9 after planting, and a scale of 1 (completely resistant) to 10 (completely susceptible) was used to rate the interaction phenotypes. Letters represent whether differences among each age group were significant (p_adj_ < 0.05). The relative order of strength observed was *Hm1*^*B73*^ > *Hm1*-*3* > *Hm1*^*A*^ > *Hm1*-*4* > *hm1*-*2*.

**Table 1.** The nature of molecular changes in the mutant alleles of *Hm1* generated by mutagenesis and their respective disease/resistance phenotypes to infection by CCR1 at maturity.

| Allele No. | Disease Response | Mutation |
| --- | --- | --- |
| *Hm1-3* | APR | T90M |
| *Hm1-4* | APR | V210M |
| *hm1-5* | Null | C82Y |
| *hm1-6* | Null | Nonsense |
| *hm1-7* | Null | Nonsense |
| *hm1-8* | Null | Nonsense |
| *hm1-9* | Null | Nonsense |

### Like *Hm1*^*A*^, the new APR alleles encode HCTRs with intermediate activity

To evaluate if the HCTR activity encoded by *Hm1-3* and *Hm1-4* was also partially compromised like that of *Hm1*^*A*^, we used the aforementioned LC-MS/MS based activity assay on samples derived from these two mutants as well as their positive and negative controls. During weeks-3 and 7 (when APR plants are susceptible and resistant, respectively), crude protein was extracted from the leaf tissue following inoculation with CCR1. The HCTR activity of extracts from APR plants was found to be significantly reduced when compared with B73 at both week-3 and week-7, indicating that HM1-3 and HM1-4 proteins display partially compromised HCTR activity during both susceptible and resistant plant ages like HM1^A^ (Fig 6). Furthermore, and consistent with HM1^A^ (Fig 3C), the levels of their HCTR did not change significantly with age (Fig 6), demonstrating that the APR encoded by these new alleles was also expressed without a concomitant increase in HCTR levels in mature plants.

**Figure 6.**
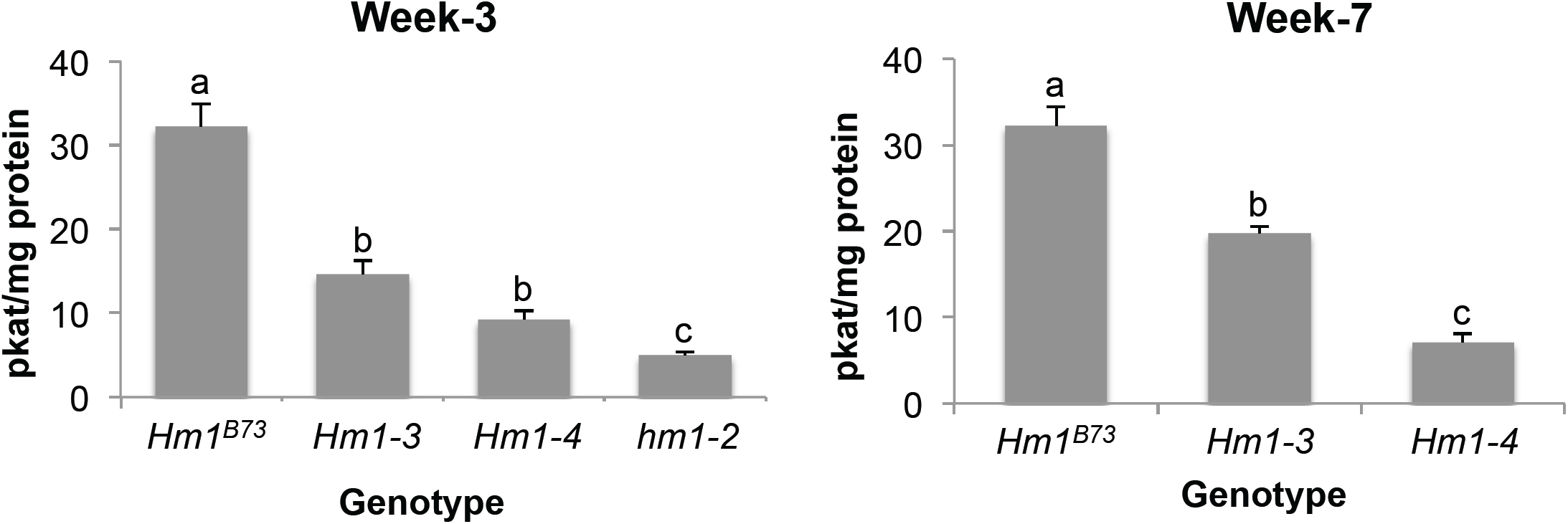
*In vitro* enzymatic activities of HCTRs encoded by the new APR alleles of *hm1*. Protein extracts from the leaf tissue of near-isogenic lines of the APR alleles *Hm1-3* and *Hm1-4* in the B73 background were used to conduct *in vitro* HCTR assays. The fully resistant (*Hm1*^*B73*^) and susceptible (*hm1-2*) alleles of *hm1* were used as controls. HCTR activities, measured at age week-3 and week-7, relied on to determining the amount of HC-toxin reduced via LC-MS/MS. Letters represent whether differences among each age group were significant (p_adj_ < 0.05).

Differences in the disease/resistance ratings of the new APR alleles predicted corresponding differences in their HCTR activities. This indeed was found to be true. The disease severity of APR plants at 3 weeks of age was found to be linearly correlated with HCTR activity (Fig 5 and 6). The APR allele with the highest degree of HCTR activity was HM1-3, followed by HM1^A^, and HM1-4 being the weakest (Fig 6). This variation in enzymatic activity is consistent with the gradient of CCR1 resistance displayed by *Hm1-3, Hm1*^*A*^, and *Hm1-4* plants from strongest to weakest (Fig 5). At maturity, however, plants carrying any of these weak alleles of *Hm1* were all indistinguishable from WT B73. This was not the case with plants carrying only the null allele; they remained uniformly susceptible to CCR1 infection even at maturity.

### Modulation of photosynthesis output alters susceptibility to CCR1 in *Hm1*^*A*^ seedlings

If the HCTR levels of the APR alleles remain largely uniform throughout plant development, why then are weak alleles unable to confer protection at the seedling stage? Some anecdotal observations that we have made about plants with APR alleles suggested that the availability of fixed carbon for energy production played a role in determining the ability of these weak alleles to suppress disease. The APR mutants always exhibited greater disease susceptibility and prolonged sensitivity in winter greenhouses as compared to the field. In the winter greenhouse, those plants closest to supplemental lights were more resistant than plants growing distant from light fixtures. Third, the resistance phenotype of APR alleles was compromised in the dominant *oil-yellow1-N1989* allele that has a chlorophyll deficiency (30).

We grew the *Hm1*^*A*^ plants at extended and reduced photoperiods to test the hypothesis that energy availability from fixed carbon could determine disease susceptibility in APR mutants. We grew *Hm1*^*A*^ B73 NIL homozygotes in a growth chamber with a light regimen of 12h light (L) and 12h dark (D) for 2 weeks. Following inoculation with CCR1 and overnight incubation, half of the seedlings were shifted to a growth chamber adjusted at 18h L and 6h D. *Hm1*^*A*^ seedlings grown in 12:12 L:D photoperiod were susceptible to CCR1 when examined at 72 hours post-inoculation (hpi) (Fig 7A) and showed no ability to suppress expanding lesions at 96 hpi (Fig 7B). However, the *Hm1*^*A*^ plants that were shifted to 18:6 L:D developed a resistant reaction instead (Fig 7C and D). Thus, the seedling susceptibility of *Hm1*^*A*^ conferred by low HCTR activity could be overcome by providing a longer period of photosynthetically active radiation.

**Figure 7.**
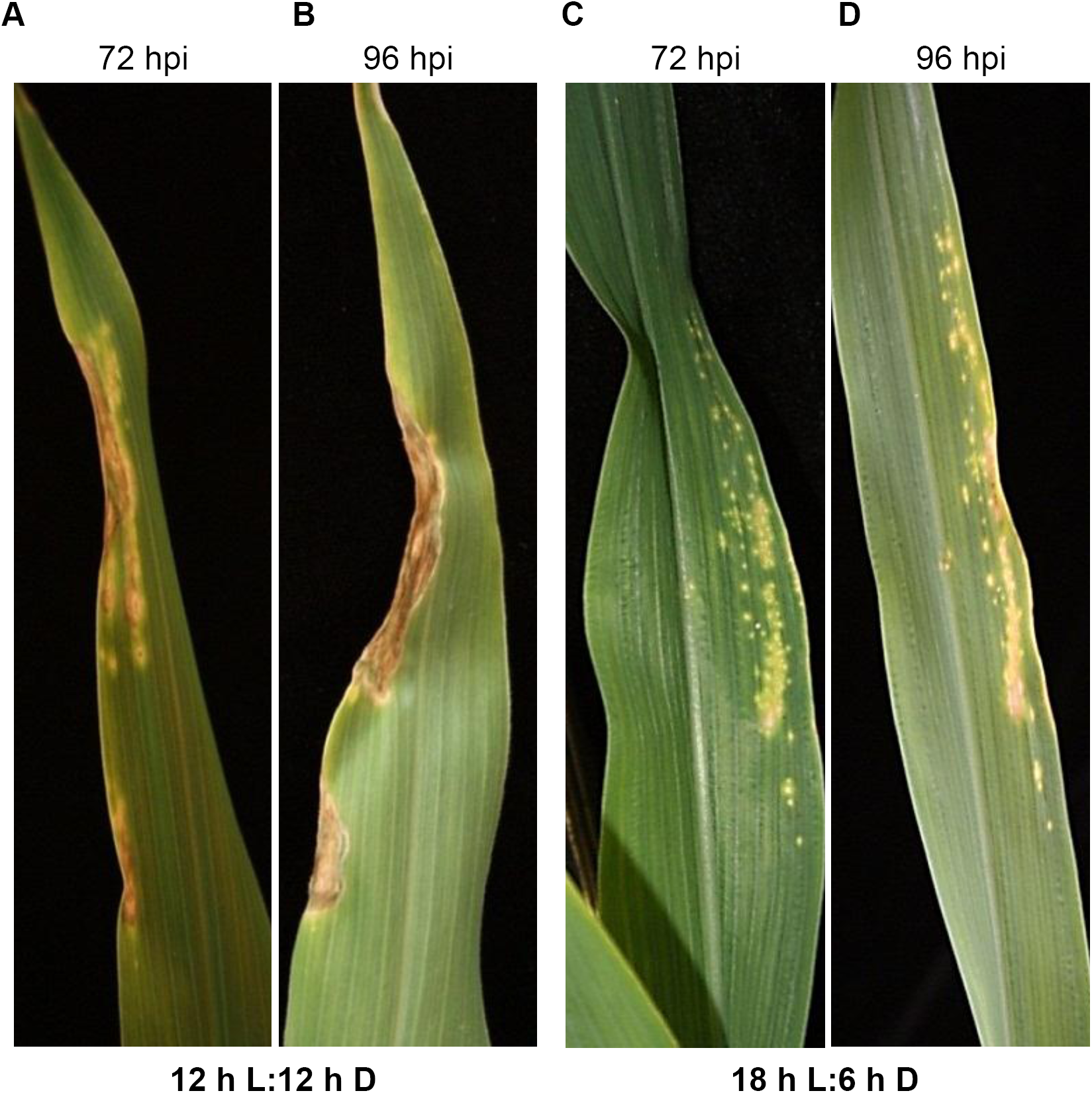
Resistance of *Hm1*^*A*^ seedlings to CCR1 in increased by extended photoperiod. Two-week-old homozygous *Hm1*^*A*^ seedlings were inoculated with CCR1 and incubated under two different photoperiods of 12 h daylight (12 h L:12 h D) and 18 h daylight (18 h L:6 h D). *Hm1*^*A*^ seedlings grown under 12 h daylight were susceptible to CCR1 at 72 hpi **(A)** and 96 hpi **(B)**. *Hm1*^*A*^ seedlings incubated under the extended photoperiod of 18 h light exhibited notably enhanced resistance at both 72 hpi **(C)** and 96 hpi **(D)**.

We reasoned that if greater photosynthate availability provides enhanced resistance sufficient to permit the weak *Hm1* alleles to confer seedling resistance, disruption of energy balance should negate their ability to confer any resistance. To test this, we treated *Hm1*^*A*^ and *Hm1-3* homozygotes with extended darkness or with the herbicide (3-(3,4-dichlorophenyl)-1,1-dimethylurea) (DCMU), which disrupts electron transfer during the light reactions of photosynthesis. We inoculated two-week-old plants with CCR1 and grew them in 14:10 L:D or 4:20 L:D. Extending the dark period of the diurnal cycle resulted in an increase in disease severity after 7 days of growth for *Hm1-3* plants (Fig 8A). If extended darkness renders plants susceptible to CCR1 due to a lack of photosynthesis, then disruption of photosynthesis by herbicide treatment should effect the same result. To test this, plants grown at 14:10 L:D were inoculated with CCR1 and grown for 24 h. At 24 hpi, plants were divided into two groups with one receiving a solution of DCMU applied to the leaf whorl and then grown for 6 days under the 14:10 L:D cycles. Observation of plants 7 dpi and 6 days after the DCMU treatment demonstrated that a single DCMU application rendered both *Hm1*^*A*^ and *Hm1-3* homozygotes completely susceptible to CCR1 (Fig 8b).

**Figure 8.**
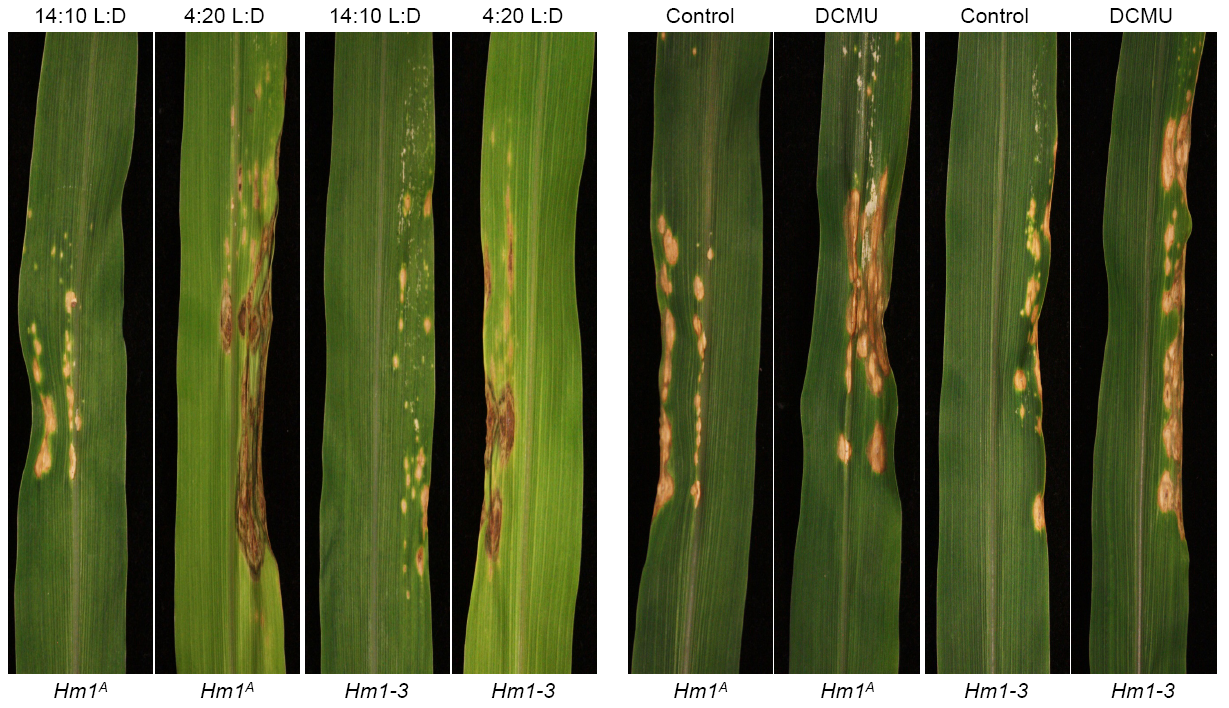
Decreased photoperiod and photosynthesis inhibition by DCMU enhanced the susceptibility of APR genotypes to CCR1. **(A)** Two-week-old homozygous *Hm1*^*Aa*^ and *Hm1-3* B73 NIL plants were inoculated with CCR1 and incubated with a shortened photoperiod of 4:20 L:D or longer 14:10 L:D photoperiod. Plants grown under a decreased photoperiod were completely susceptible to CCR1 while control plants were relatively less susceptible. **(B)** *Hm1*^*A*^ and *Hm1-3* B73 NIL plants were grown for two-weeks in the longer photoperiod conditions (14:10 L:D) and half of the plants were sprayed with DCMU, a photosynthesis inhibiting herbicide. Application of DCMU rendered both *Hm1*^*A*^ and *Hm1-3* plants highly susceptible to CCR1 compared to control plants. Pictures were taken 6 days after inoculation.

Together, these two experiments demonstrate that light, and perhaps the energy status of the plant, were key determinants of resistance to CCR1, and provide a direct link between plant primary metabolism and physiology and disease resistance.

## DISCUSSION

This study reveals one fundamental aspect of adult plant resistance (APR) in maize to CCR1. APR alleles at the *hm1* locus are weak determinants of resistance that fail to protect plants at the seedling stage but are sufficient to confer complete protection to CCR1 at maturity. This conclusion is supported by multiple lines of evidence derived from a combination of genetic, molecular, and biochemical experimentation. Genetic analysis demonstrated that all APR alleles of *hm1* confer partial resistance that exhibits haploinsufficiency (gene-dosage sensitivity) during most stages of plant development. This contrasts with resistance conferred by the wild type (WT) alleles of *hm1* that are completely dominant and protect every part of the plant regardless of age or maturity. Plants with null alleles of *hm1,* on the other hand, are susceptible to CCR1 at all stages of development. CCR1 infection typically results in plant lethality for these alleles, and the ubiquitous nature of this pathogen makes them difficult to propagate in the field. The APR alleles of *hm1* are recessive to the WT alleles (e.g., *Hm1*^*B73*^) but dominant to null alleles of *hm1* (e.g., *hm1*^*Pr*^).

Consistent with the idea that APR is a symptom of weak or partial loss-of-function alleles, we were able to generate two new APR alleles from the WT *Hm1*^*B73*^ allele by mutagenesis with EMS. Five completely susceptible mutants were also recovered in this mutant screen, which presumably encoded null mutations. In keeping with these predictions, molecular analysis of these null alleles showed that four of the five null mutants were the result of nonsense mutations that truncated their predicted peptides by introducing premature stop codons. The fifth null mutant, which was caused by a missense mutation, changed a highly conserved cysteine residue (C82Y) that is perhaps critical for protein function. In sharp contrast, both novel APR alleles underwent relatively conservative mutational changes: T90M in *Hm1-3* and V210M in *Hm1-4*. Even *Hm1*^*A*^, which differs from the WT *Hm1*^*B73*^ allele by five amino acids, seems to owe its APR phenotype to a single L116H change. HCTR activity was encoded by all of the APR alleles, indicating that none of these mutations completely eliminates the function of the enzyme. Their HCTR activities were compromised, however, being intermediate to that of the fully functional WT allele (which confers completely dominant protection) and the recessive null *hm1* alleles, which impart no resistance to CCR1. These results indicate that at some level HCTR activity is unable to deter the pathogen from colonizing maize plants at the seedling stage but that level of activity is sufficient to prevent CCR1 from colonizing at maturity.

A cause-and-effect relationship between APR and partial-loss-of-function alleles of *hm1* is further substantiated by the correlation between the strength of the resistance reaction conferred by an APR allele and its HCTR activity. The level of HCTR activity matched perfectly with the strength of CCR1 resistance conditioned by the three APR alleles. These results demonstrate that alleles of *hm1* with partial loss-of-function mutations encode HCTR with a compromised activity and that the weaker activity results in later onset of disease resistance. The resistance of seedlings encoding WT *Hm1* demonstrates that efficient toxin deactivation is sufficient for maize seedlings to resist CCR1 infection and, therefore, they express all of the required machinery for defense. Likewise, mature plants lacking *hm1* function are completely susceptible, demonstrating that HCTR is absolutely required for CCR1 infection, and mature maize plants are not protected from toxin-mediated disease spread. These interpretations depend on the *in vitro* assay correctly reflecting *in vivo* activity. Our *in vitro* HCTR activity assay did not detect the *in vivo* activity of the enzyme but instead the level of the functional protein present at a given time point. It is possible that *in vivo* activity did not correspond to the *in vitro* activity identified by this method.

A seemingly mechanistic relationship between partial resistance and APR is also evident in many other pathosystems where such genes have been cloned and studied in detail. One example is that of *Cf-9B*, which mediates incomplete resistance to *C. fulvum* in a developmentally specified fashion (17). Its paralog *Cf9*, which encodes a receptor like protein, confers complete protection in all plant tissues at every stage of development (31). Another example is that of *Xa21*, a receptor-like kinase that confers weak resistance to Xanthomonas leaf blight in rice (14,15). The maize *Hm2* APR allele provides another example. The weak CCR1 resistance provided by this allele is conferred by a truncated HCTR (12).

In wheat, APR genes are rather common and have been used widely to protect this crop from all forms of the disease caused by three different species of rust pathogens (reviewed in Ellis et al., 2014). Even though APR genes confer little or no protection in wheat seedlings, the broad-spectrum and durable nature of resistance provided by such genes in adult plants have many breeders proclaim that breeding for rust resistance should deploy only APR genes (32). Three of these wheat APR genes have been cloned recently and, interestingly, they all appear to confer resistance by different mechanisms. One of them, *Yr36*, a mediator of resistance to yellow rust, encodes a kinase with an unusual domain (16), while *Lr34* and *Lr67*, both of which mediate APR to both rust and powdery mildew pathogens, encode an ABC transporter and a hexose transporter, respectively (13,19). Exactly how these genes confer APR remains unresolved, but one thread that unifies them is their ability to confer only weak or partial resistance (32). Overexpression of *Lr34*, one of the best studied APR genes, however, did enable it to confer seedling resistance in durum wheat (33). Furthermore, the efficacy of this transgene in conferring seedling resistance improved even further under extended daylight conditions (34). These results echo what we have discovered with the APR alleles in maize and suggest that the connection between weak resistance and APR is not unique to the maize-CCR1 pathosystem but perhaps is a general feature of most disease resistance genes that are weak and provide only partial protection.

A second major finding is that APR is not the result of the enhanced level or activity of proteins encoded by APR alleles at the mature-plant stage. Rather, it must be the result of a change in seedlings vs mature plants that affects differential resistance. It was previously shown in a number of cases that the differential transcriptional activity of an APR gene did not account for its APR phenotype (12,13,15,17,20). Here we extend this to the level and HCTR activity of the accumulated HM1 proteins, which remained stable across development. At the onset of APR, resistance manifests uniformly in all parts of the plant, including the youngest leaves that are still unfurled, indicating that the APR-inducing factor is not accumulated over a long period of time in aging tissues, but rather is available in every part of the plant regardless of the age of the organ and determined solely by the plant maturity.

Considering that the HCTR activity is present at equivalent levels in APR mutant extracts regardless of plant stage, why then are seedlings susceptible? Though the studies presented here do not resolve this question, the biochemical mechanism by which *hm1* confers resistance to CCR1 suggests a plausible scenario. Although this resistance is conferred by *hm1*-encoded HCTR, the HC-toxin (HCT) inactivation reaction requires the reducing power of NADPH as a co-substrate. The direct involvement of NADPH in HC-toxin reduction suggests this molecule could be very critical in regulating resistance in the maize-CCR1 pathosystem. Supporting this hypothesis are our results showing that light and photosynthetic activity have a great impact on resistance mediated by APR alleles, either boosting them to confer seedling resistance or limiting them to prevent APR.

Based on these results, it could well be the availability of NADPH that determines the difference in resistance between seedling and mature stages in the *hm1* APR mutants. NADPH is produced during the light reactions of photosynthesis, the C4 malate shuttle, and sugar oxidation, along with other energy carriers such as ATP. Maize seedlings not only have a limited photosynthetic capacity to assimilate carbon (C), but also strong sinks to consume these assimilates (35). As a result, seedling leaves become C-deficient at night and that may negatively impact the availability of NADPH and ATP. Since NADPH is required for HCTR activity, its depletion at night may negatively impact the activity of hypomorphic mutants of HCTR, thereby leaving HCT active to induce susceptibility to CCR1. Bolstering this hypothesis is the observation that the *Hm1-3* and *Hm1-4* mutations occur at residues predicted to be critical for the binding of NADPH to HCTR (36). The WT HCTR has likely evolved to require lower NADPH levels for optimal activity, buffering any impact from the likely diurnal dip in its cofactor at night and thereby allowing sufficient HCT inactivation. This scenario also explains why plants with the APR genes become more resistant as they mature; the increased output of photosynthates may outstrip the sink requirements, allowing excess photosynthates to be stored as starch during the day and then used at night to fuel NADPH production.

Although several other aspects of plant bioenergetics are expected to support the resistance phenotype of the APR genes in most pathosystems, NADPH appears to be the most critical in energizing APR in the maize-CCR1 pathosystem. This, of course, is due to the direct involvement of this molecule in the resistance mechanism mediated by HCTR, and is supported by the fact that maize plants carrying the WT *Hm1* gene are completely resistant to CCR1 at all stages of development, including as seedlings. This study thus provides direct evidence linking, for the first time, primary host metabolism to the realm of disease resistance in plants.

An intriguing implication of this study concerns the metabolic cost of resistance in plants. This topic is not only of fundamental interest to plant pathologists and entomologists but also has huge agricultural relevance (37–39). Our study demonstrates that, compared to strong resistance, the weak form of resistance has a much higher metabolic cost for the host. As shown in the case of APR, this cost can be so high that the seedlings are not robust enough metabolically to express such resistance effectively. This argument also extends to the quantitative form of resistance that is often relatively weak and easily affected by the environment (40,41). An additional complication is that the vulnerability of seedlings to diseases increases even further by conditions that compromise photosynthesis. This phenomenon is analogous to what has been well established in the animal world that malnutrition compromises the immunity of infants much more than that of adults (42,43).

## METHODS

### Plant materials

The inbred P8 and landraces Pira and Enano were obtained from Germplasm Resources Information Network (GRIN) of the U.S. National Plant Germplasm System. The CCR1-susceptible maize inbred Pr, and the CCR1-resistant inbreds B73, Va35, W22, and Pr1 (a near-isogenic line of Pr) were previously available in our research program. To determine whether *Hm1*^*A*^ is an allele of *Hm1*, P8 was crossed with Pr and the F_1_ hybrid was backcrossed to Pr to generate a BC_1_F_1_ population. Additionally, P8 was crossed with Pr1 and the resulting F_1_ hybrid was testcrossed to the *hm1* null stock Pr. Near-isogenic lines of B73 displaying APR to CCR1 infection were generated by backcrossing *hm1* APR alleles with the B73 inbred, to determine the behavior of the APR alleles in a uniform genetic background.

### Pathogen growth and inoculation

The protocol for culturing CCR1 pathogen on carrot juice agar medium was the same as previously described (23). One-hundred μl of 10^5^ spores/ml of CCR1 conidial suspension was used for leaf whorl inoculations. To study the phenotypic manifestation of APR by the *Hm1*^*A*^ allele, both homozygous (*Hm1*^*A*^*Hm1*^*A*^ introgressed into B73) and heterozygous (*Hm1*^*A*^*hm1*^*Pr*^ also in B73) plants were planted in isolation at the Purdue ACRE farm and inoculated with 100 μl of 10^5^ spores/mL of CCR1 spore suspension. Wild type B73 encoding *Hm1*^*B73*^ and the susceptible *hm1*^*Pr*^ B73 NIL plants were used as resistant and susceptible controls, respectively. A fresh set of five rows of ∼40 plants per row was inoculated every week, and disease severity rating was determined 5 days post-inoculation (dpi) as described previously (12). To determine if *Hm1*^*A*^ is an allele of *Hm1*, genetic crosses were made at the ACRE farm and the resulting segregating progeny was evaluated under field conditions again at the ACRE farm.

### Amplification of *Hm1*^*A*^ genomic DNA

Four primer pairs were designed to amplify *Hm1*^*A*^ based on its sequence homology with *Hm1*^*B73*^. The promoter region was amplified using a primer pair based on the promoter of *hm1* from Pr. Touchdown PCR (44) was carried out with 10 consecutive cycles of denaturation at 94°C for 30 sec, annealing at 63°C for 30 sec with a decrease in 0.5°C per cycle to a “touchdown” of 58°C, and extension at 72°C for 30 sec; followed by 35 cycles of 94°C for 30 sec, 58°C for 30 sec, and 72°C for 45 sec. Three separate PCR reactions were carried out for every primer so that any errors initiated by either the GoTaq DNA Polymerase (Promega, Madison, WI, USA) or by sequencing could be ruled out. The PCR products were cleaned by running them through an agarose column, BigDye sequencing reactions were conducted, and the products precipitated with sodium acetate and ethanol before final resuspension in 20 μl of double-distilled water (ddH_2_O). These samples were submitted to the Purdue Genomics Facility for low throughput sequencing. Forward and reverse complementary sequences for each primer were compared using the ClustalW2 multiple alignment program. In order to assemble the *Hm1*^*A*^ sequence without sequencing errors, only sequences with at least three perfect reads for each primer sequence were considered.

### Cloning of *Hm1*^*A*^ cDNA

P8 (*Hm1*^*A*^*Hm1*^*A*^) seeds were planted in 500M MetroMix and grown in Conviron growth chambers for two weeks. One-hundred μl of 10^5^ spores/mL CCR1 spore suspension was used for whorl inoculation, and plants were covered with a hood overnight to maintain humidity required for spore germination and penetration into the leaf tissue. At 24 h post-inoculation (hpi), affected leaf tissue was collected from the plants and snap-frozen in liquid nitrogen. RNA was extracted with a Qiagen RNeasy extraction kit (Qiagen, Germantown, MD), and cDNA was synthesized by RT-PCR using random hexamer mix (New England BioLabs, Ipswich, MA).

### Generating near-isogenic lines of B73 manifesting APR and susceptibility to CCR1

The P8 maize inbred line was crossed with the maize reference B73 inbred, and the resulting F_1_ hybrid was backcrossed to B73. To introgress *Hm1*^*A*^ into the B73 inbred, the resulting BC_1_F_1_ progeny was backcrossed to B73 for six generations. Since the promotor region of *Hm1*^B73^ differed from that of *Hm1*^*A*^, PCR-based markers designed from the promotor region were used for introgressing *Hm1*^*A*^ into B73 (primer sequences are available in Table S1). After the BC_7_ generation, *Hm1*^*A*^ containing plants (*Hm1Hm1*^*A*^) were self-pollinated to generate homozygous *Hm1*^*A*^ B73 NIL plants. Homozygous *Hm1*^*A*^ B73 NIL plants were identified with PCR-based markers and were self-pollinated to generate seed. Similar to *Hm1*^*A*^, the two novel APR alleles *Hm1-3* and *-4* generated through EMS mutagenesis were introgressed into the B73 inbred for seven generations using a Cleaved Amplified Polymorphic sequences (CAPs) assay (primer sequences in Table S1). The restriction enzyme *NlaIII* (New England BioLabs, Ipswich, MA) was used to differentiate the *Hm1*^*B73*^ allele from the two novel APR alleles. Similar to the novel APR alleles, the novel null allele *hm1-2* identified in the EMS-mutagenized B73 M_2_ family screen was backcrossed for five generations into B73 using PCR-based markers and self-pollinated to obtain a homozygous *hm1-2* NIL in B73. Marker-assisted backcrossing using PCR-based genotyping was conducted on plants grown at the Purdue Agronomy Center for Research and Education (ACRE) farm during the summer and in the Purdue University Botany and Plant Pathology greenhouses during the winter season.

### Transcriptional activity of *Hm1*^*A*^

*Hm1*^*A*^ plants were inoculated with CCR1 spore suspension as described above at weekly intervals from the seedling stage to maturity (week-1 through week-8). Total RNA was isolated from CCR1-infected leaf tissue as described by Eggermont et al. (1996) and treated with RNase-free DNase I to eliminate genomic DNA using the TURBO DNA-free Kit (Ambion, Austin, TX). One μg of treated RNA was reverse-transcribed to cDNA in a total volume of 25 μl using the iScript^TM^ cDNA Synthesis kit from Bio-Rad (Hercules, CA). RT-PCR was conducted using gene specific primers with the maize actin transcript as a control (see Table S1 for primer information). RT-PCR was conducted under the following conditions: denaturation at 94°C for 1 min, annealing at 60°C for 1 min, extension at 72°C for 1 min, and a terminal extension steps for 10 min. 30 and 28 cycles of PCR were conducted to amplify *Hm1*^*A*^ and the control actin gene, respectively. Amplified PCR products were separated on a 0.8% agarose gel to visualize the expression of the *Hm1*^*A*^ transcript. Three replicates for each time point were used for this experiment.

Additionally, qRT-PCR was conducted on cDNA from *Hm1*^*A*^ plants inoculated with CCR1 at week-1, −3, −5, and −7 using gene specific primers. For relative quantification, Molybdenum co-factor biosynthesis protein (MOL, GRMZM2G067176) was used as a reference gene (46). All primer combinations had an efficiency of 90-100%. Individual qRT-PCR reactions contained 5 μl of SYBR^®^ Select Master Mix (Applied Biosystems, Foster City, CA), 2 μl of cDNA template (20x dilution), and the appropriate amount of forward and reverse primers plus water. A three-step qRT-PCR amplification (40 cycles of 95°C for 5 s followed by 61°C for 20 s and 72°C for 30 s) was performed using the Mx3000P qPCR system (Stratagene–Agilent Technologies, Santa Clara, CA). Semi-quantitative RT-PCR was conducted using gene-specific primers for *Hm1*^*A*^ and the reference gene *Actin* (primer sequence in Table S5). Three replicates for each time point were used for this experiment.

### Translational activity of *Hm1*^*A*^

For Western analysis, samples were collected at weekly intervals from week 2 through week 8 after planting. Since *Hm1*^*A*^ is infection inducible, plants were inoculated with CCR1 for 24 h before collecting leaf samples. Protein was extracted using an alkaline lysis protocol and quantified using the Bradford method. Equal amounts of the protein (10 ug) was loaded onto SDS-PAGE gels and western blots were developed using a MAP antibody raised against a synthesized 13 amino acid peptide (RPARDRLGELGFK) corresponding to amino acids 312 to 324 of the HM1 peptide.

For the HCTR activity assay, *Hm1*^*B73*^, *hm1*^*Pr*^, and *Hm1*^*A*^ plants grown in the field were inoculated with 200 μl of 10^5^ spores/mL CCR1 spore suspension into the leaf whorl at weeks-3 and −7. Four biological replicates of three inoculated plants were sampled 24 hpi and stored at −80°C until used. Total plant protein was extracted using a protocol adapted from Hayashi et al. (2005) and desalted using a Sephadex G-50 Fine column (GE Healthcare, Chicago, IL). After determining protein concentration with a Bradford assay, 13.55 μg of protein was used to start reactions containing 25 mM Tris-HCl (pH 7.0), 160 mM NADPH, and 55 μM HC-toxin. The assays were run at 30°C in the dark for 45 min and then stopped by the addition of 1.25 ml cold acetone. After centrifugation at 15,000 x *g* for 15 min at 4°C, 10 μL of the supernatant was injected onto an Atlantis T3 column (2.1 x 150 mm, 3 μm, 100 Å, Waters) maintained at room temperature and analyzed using an Agilent 1200 series LC instrument coupled to an Agilent 6460 triple quadrupole mass spectrometer (Agilent Technologies, Santa Clara, CA) at the Bindley Bioscience Center in Purdue Discovery Park.

The solvent system contained solvents A (0.1% formic acid in ddH_2_O) and B (0.1% acetonitrile). The column was eluted with 85% A and 15% B (0 to 1 min), followed by a linear gradient from 1 to 16 min to 40% A and 60% B, and a hold from 16 to 16.5 min at 40% A and 60% B. The column solvent was then reduced from 60% B to 15% B (16.5 to 17 min) and kept isocratic at 15% B from 17 to 22 min with a flow rate of 0.3 ml/min. HC-toxin (Sigma-Aldrich, St. Louis, MO) and its reduced form eluted from the column at 8.5–11.5 min under these conditions. During the analysis, the column effluent was directed to the MS/MS, with the Jetstream ESI set to positive mode with nozzle and capillary voltages at 1000 – 4000 V. The nebulizer pressure was set at 35 psi, the nitrogen drying gas was set at 325°C with a flow rate of 8 L/min, and the sheath gas was held at 250°C at a flow rate of 7 L/min. Fragmentation was achieved with 70 V for both analytes. Multiple reaction monitoring (MRM) was used to selectively detect HC-toxin and its reduced form. The first quadrupole was set to transition between the [M-H]^+^ of the analytes, whereas the last quadrupole monitored *m/z* 411 and 409 for reduced and normal HC-toxin respectively. Each transition was monitored with a dwell time of 150 ms and collision energy of 15 V, with ultrapure nitrogen used as the collision gas. Mass selection was achieved using the following ions: 439.3 for reduced HC-toxin and 437.3 for HC-toxin. Data were collected and analyzed via the MassHunter Workstation (version B.06.00, Agilent Technologies, Santa Clara, CA), and peak areas were determined by integration. Similar to *Hm1*^*A*^, the HCTR activity of the new APR alleles generated by targeted EMS mutagenesis (*Hm1-3* and *Hm1-4*) along with resistant (*Hm1*) and susceptible (*hm1-2*) controls were also evaluated by LC-MS/MS.

### Generating novel APR manifesting alleles of *Hm1* by EMS mutagenesis

The B73 (*Hm1Hm1hm2hm2*) maize inbred, which exhibits complete resistance to CCR1 at all stages of plant development (23), was the pollen parent for the targeted EMS mutagenesis screen. The CCR1-susceptible maize inbred Pr (*hm1hm1hm2hm2*) (9,24), which exhibited complete susceptibility to CCR1 at all stages of plant development, was used as the female parent. This experiment was conducted in a greenhouse facility, as the Pr plants do not survive in the field due to high levels of disease pressure.

To conduct pollen EMS mutagenesis, EMS stock solution was prepared by adding 1 ml of EMS (Sigma-Aldrich, St. Louis, MO) to 99 ml of paraffin oil (Sigma-Aldrich, St. Louis, MO). Tassels of the Pr plants were removed before starting the experiment. On the day of conducting pollen mutagenesis, EMS working solution was prepared by mixing 1 ml of EMS stock solution with 14 ml of paraffin oil. This working solution of EMS was mixed gently for one hour to uniformly disperse the EMS in paraffin oil. B73 pollen was collected in tassel bags, measured and transferred to a 50-ml Nalgene bottle. For every 1 ml of pollen collected, 10 ml of EMS working solution was added. The EMS-treated pollen was placed on ice and mixed gently every 5 min for 45 min. About two to three drops of EMS-treated B73 pollen was then applied to the silks of Pr ears. Ears from these Pr plants were harvested 45 days after pollination. The M_1_ seeds (∼4500) obtained from this genetic cross were planted at the Purdue ACRE farm. At both week-2 and week-5, plants were whorl-inoculated with 100 μl of 10^5^ spores/mL of CCR1 conidial suspension and screened for their disease response one week post-inoculation.

### Amplification of *Hm1*^B73^ allele from heterozygous CCR1-susceptible mutants

Based on sequence polymorphisms between the wild type *Hm1* from B73, *Hm1*^*B73*^ and the null *hm1* allele from Pr, *hm1*^*Pr*^, four primer pairs amplifying −560-bp of the promoter region from the translation start site and the entire coding region of *Hm1* were designed to preferentially amplify the WT *Hm1*^*B73*^ from heterozygous M_1_ plants (Fig S5), which were obtained by crossing Pr plants with EMS-treated B73 pollen. Four overlapping primer combinations (primer sequences in Table S1) were used to preferentially amplify *Hm1*^*B73*^ over the *hm1*^*Pr*^ allele. Amplified PCR fragments were processed as described above for *Hm1*^*A*^ amplification and submitted to the Purdue Genomics Facility for low-throughput sequencing.

### Differential photoperiod treatments of *Hm1*^*A*^ plants

*Hm1*^*A*^ B73 NIL plants were grown in Conviron growth chambers providing a 12:12 L:D photoperiod. Two-week-old *Hm1*^*A*^ plants were inoculated with 100 μl of 10^5^ spores/mL of CCR1 spore suspension into the leaf whorl. CCR1-inoculated plants were incubated overnight in a humidity chamber at 80% relative humidity. These plants were then subjected to 12:12 L:D or 18:6 L:D photoperiods. The response reaction to CCR1 infection was evaluated every 24 h for a 96 h period. Digital photographs of lesion progression were taken using a Canon EOS Digital Rebel XSi camera.

Additional extended darkness and DCMU treatment experiments were performed in growth chambers on plants homozygous *Hm1*^*A*^ and *Hm1-3* in the B73 genetic background. Plants were grown in a growth chamber under 14:10 L:D for two weeks. We inoculated these plants with CCR1 and subjected them to two different light regimes, 14:10 L:D or 4:20 L:D. On a subset of CCR1 inoculated plants transferred to 14:10 L:D, the herbicide (3-(3,4-dichlorophenyl)-1,1-dimethylurea) (DCMU) at a concentration of 100 μM was applied to the leaf whorl 24 hpi. Disease severity of these plants was determined at 7 dpi.

## ACKNOWLEDGEMENTS

This work was partially supported by GSJ’s Hatch project (IND011280), by the IOS-NSF grant 0547132 to G.S.J., and by the National Science Foundation Plant Genome Research Program grant 1444503 to B.P.D. and G.S.J.

## AUTHOR CONTRIBUTIONS

DSM, SC, SM and GSJ designed and performed linkage analysis studies. DSM did the Southern blots and also constructed and screened the lambda library to clone *Hm1*^*A*^. SM conducted sequence alignment analysis with *Hm1* orthologs and different maize inbreds. SC performed *Hm1*^*A*^ transcriptome analysis using semi-quantitative RT-PCR and AK conducted qRT-PCR. KC and AK designed and performed HCTR assays. SC, SM, and GJ designed and conducted EMS mutagenesis. SM, KC, and GJ screened EMS-generated M_1_ plants for their disease response to CCR1. SM, AD, BK, BD and GJ designed and performed experiments to look at the effect of photosynthesis on APR. LD initially provided P8, the inbred possessing *Hm1* A, and worked with GJ to characterize its APR. SM, SC, KC, BD, and GJ wrote the paper. All authors reviewed the manuscript and LD provided the detailed editorial changes.

